# Discovery and reporting of clinically-relevant germline variants in advanced cancer patients assessed using whole-exome sequencing

**DOI:** 10.1101/112672

**Authors:** Tuo Zhang, Alessandro Romanel, Kenneth W. Eng, Hanna Rennert, Adrian Y. Tan, Yaohua Xue, Joanna Cyrta, Juan Miguel Mosquera, Andrea Sboner, Ivan Iossifov, Steven M. Lipkin, Jenny Xiang, Xiaojun Feng, Peter Nelson, Himisha Beltran, Colin C. Pritchard, Mark A. Rubin, Francesca Demichelis, Olivier Elemento

## Abstract

**Purpose:** In precision cancer care, WES-based analysis of tumor-normal samples helps reveal somatic alterations but can also identify cancer-associated germline variants important for disease surveillance, treatment choice and cancer prevention. WES can also identify germline secondary findings impacting risk of cardiac, neurodegenerative or metabolic diseases. In patients with advanced cancer, the frequency of reportable secondary findings encountered with WES is not well defined.

**Methods:** To address this question, we analyzed a cohort of 343 patients with advanced, metastatic cancer for whom we have performed tumor and germline WES interrogating more than 21,000 genes using a CLIA/CLEP approved assay.

**Results:** 17% of patients in our cohort have one or more reportable germline variants, including patients with pathogenic variants in the BRCA1 and BRCA2 genes. The frequency of non-cancer clinically relevant germline variants (8.8%) was within the range of two control non-cancer cohorts (11.0% and 6.5%). The frequency of variants in cancer-associated genes was significantly higher (p<0.0005) in our advanced cancer cohort (8.2%) compared to control cohorts (2.7% and 3.8%). More than 50% of patients with reportable germline cancer variants had a family history of cancer.

**Conclusion:** these results stress the importance of returning germline results found during somatic genomic tumor testing.

## Introduction

Clinical whole exome sequencing (WES) is increasingly used in the constitutional setting for improving diagnosis of rare diseases ^1^. In the oncology setting, most institutions favor targeted sequencing panels that examine from 40 to 400 genes within tumor samples. Nonetheless a limited number of institutions have deployed whole exome sequencing in the clinic ^2–4^. Unlike with targeted panels, clinical WES typically involves sequencing both a tumor sample and a matched germline sample, then identifying somatic alterations, that is, variants that are specific to a tumor sample and not found in the matched germline sample. The systematic sequencing of a germline sample together with broad coverage provided by WES provides an opportunity to examine germline variants that may be of clinical importance for patients. Such variants fall in two categories: (1) cancer-associated variants that may predispose patients for certain types of cancers and in some cases may help identify additional treatment options and (2) non-cancer variants that may potentially impact risk of developing cardiac, neurodegenerative or metabolic diseases.

In the context of genomic tests whose primary purpose is to identify somatic mutations within tumor samples, these two categories of germline alterations are typically referred as secondary findings. Clinically relevant germline variants typically occur in specific genes such as those defined and published by the American College of Medical Genetics and Genomics (ACMG)'s Working Group on Incidental Findings in Clinical Exome and Genome Sequencing ^5^. Additional diseases-specific genes may be found in panels such as the BROCA panel for cancer risk genes ^6^ and in established pharmacogenomics databases such as PharmGKB ^7^. The list of reportable genes is continuously evolving as new clinical evidence is uncovered ^8^.

In the oncology setting there are compelling clinical reasons to report relevant secondary germline variants uncovered during the process of somatic genomic testing. For example pathogenic variants in constitutional risk genes such as BRCA1, TP53, ATM and many others are associated with increased risk of developing certain tumors. Patients may not be aware of the presence of such pathogenic variants in their genome prior to testing. Such variants may also have been inherited by siblings and shared in other family members who may want to know about such findings and take appropriate preventive action when possible. In some cases, the detection of germline alterations may not only affect family risk but also choice of systemic therapy for an advanced cancer patient, e.g., PARP inhibitor treatment in patients with germline BRCA variants. Non-cancer associated variants that increase risk of cardiac, neurodegenerative or metabolic diseases also fall in the reportable category ^5^.

Several studies have started to explore the frequency of reportable incidental germline variants uncovered by targeted, whole exome and whole genome sequencing. Many of these studies were performed in non-oncology setting ^9–12^. For example, in a WES germline cohort of patients with Mendelian disorders with unknown molecular basis, 0.86% of individuals had a reportable secondary variant among the 56 recommended genes by the American College of Medical Genetics (ACMG) ^13^. Perhaps more relevant to the present study, several studies have now reported elevated number of germline cancer predisposing variants in cancer patients. In a study using whole-genome and whole-exome sequencing involving 1,120 children and adolescents with cancer, 8.5% of the patients were found to have cancer-predisposing gene mutations ^14^. In another recent study, the incidence of germline variants in genes mediating DNA-repair processes among men with metastatic prostate cancer was found to be 11.8% ^6^. These studies have focused on specific patient populations, e.g., non-cancer research cohorts, pediatric patients and metastatic prostate cancer patients and may not represent the broader patient population encountered in routine clinical care where whole exome sequencing is clinically compelling such as advanced cancer patients.

To investigate the frequency of reportable incidental findings in a heterogeneous advanced cancer cohort, we analyzed a cohort of 343 patients with advanced, metastatic cancer for whom we have performed tumor and germline WES interrogating more than 21,000 genes using a CLIA/CLEP approved assay ^15,16^. We have recently described the landscape of somatic alterations (point mutations, indels, copy-number alterations) in this cohort ^15,16^. We describe here a comprehensive analysis of secondary germline findings in that same cohort.

## Results

### Overview of the Englander Institute for Precision Medicine (IPM) cohort, germline variant detection and ethnicity inference

We sequenced 345 germline DNA samples from 343 prospectively enrolled patients with advanced cancer as part of an IRB approved protocol (IRB #1305013903) with informed consent at the Englander Institute for Precision Medicine (IPM) of Weill Cornell Medicine/New York Presbyterian Hospital. The most common cancer types in our cohort are metastatic prostate cancer (n=97 patients), bladder cancer (n=42) and central nervous system (CNS) tumors (n=41), followed by kidney cancer (n=32), non-small cell lung cancer (n=17), breast cancer (n=12), hematological malignancies including leukemias and lymphomas (n=12), colon cancer (n=11), and ovarian cancer (n=8). These nine cancer types cover about 80% of patients in the advanced cancer cohort (Fig 1a). Samples were sequenced at an average depth of 83X using the Agilent HaloPlex whole-exome sequencing (WES) platform ^15,16^. We implemented a germline variant calling pipeline based on the BWA aligner ^17^, and GATK for base recalibration, realignment around indels and variant calling ^18^. Overall we identified 26,473 variants on average per individual. To evaluate the accuracy of the HaloPlex platform together with our variant calling approach, we sequenced and analyzed DNA from the reference NA12878 sample and compared variant calls using our approach with published benchmark reference variant calls ^19^. We tested our pipeline on two replicate NA12878 samples. The pipeline called 17,115 and 17,258 variants in the two replicates, respectively, out of which 143 and 154 were in disagreement with the reference calls (average false discovery rate 0.86%). The pipeline also missed 802 and 743 variants present in the reference dataset. Altogether, this indicates an average sensitivity rate of 95.6% and an average specificity rate > 99.999%. Two germline samples were also sequenced twice. The overall agreement between these duplicate samples was 95% on average (**Supplementary Table S1**). Most importantly, all clinically relevant reportable germline variants were identified in both replicates.

**Figure 1:**
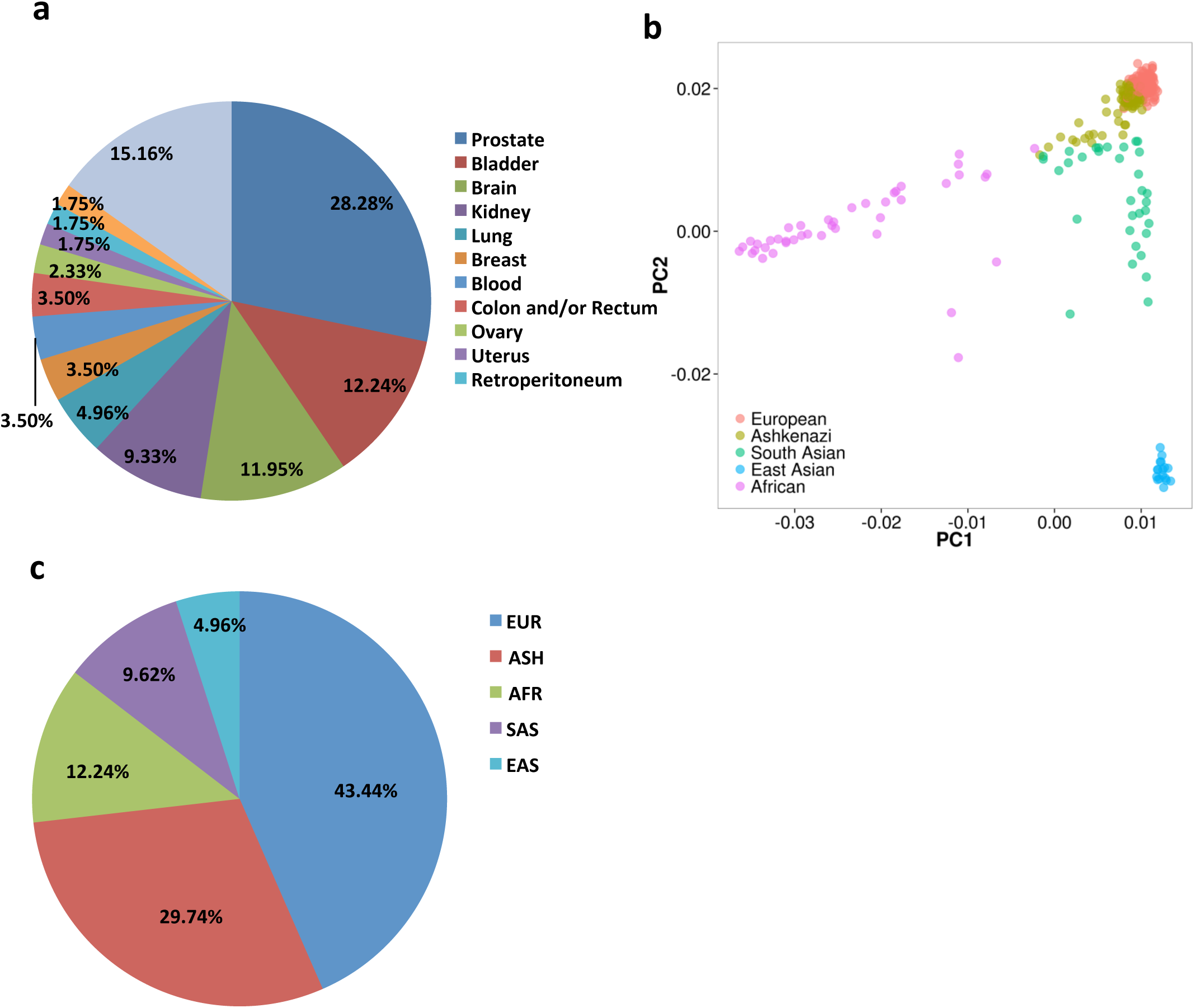
Overview on IPM cohort. (a) Cancer diagnosis in our 343 patients cohort. (b) Ethnicity map using principal component analysis, based on 35,039 SNPs. (c) Ethnicity composition inferred by Ethseq (Romanel et al, submitted), EUR for European; ASH for Ashkenazi; AFR for African; SAS for South Asian; EAS for East Asian.

To avoid well-known accuracy problems with self-reported ethnicity, we sought to infer ethnicity based on single nucleotide polymorphism (SNP) genotype calls. We used a computational approach (EthSeq) based on differential SNP-profiles that infers individual's ethnicity relying on the major ethnic groups represented in the 1000 Genomes Project (Romanel *et al,* submitted; see **Methods**). A principal component analysis (PCA) using the first two components identified three major groups in our advanced cancer cohort (Caucasians-Ashkenazim, Asians and Africans; Fig 1b) and further revealed the ethnic breakdown as 43% European Caucasian, 30% Ashkenazim and 12% Africans (Fig 1c). This analysis reveals a relatively high prevalence of Ashkenazi individuals in our New York City cohort. In comparison, a TCGA cohort with 434 blood samples contained approximately 3.5% Ashkenazim when analyzed using the same approach and parameters (data not shown).

### Variant prioritization and classification scheme reveals pathogenic or likely pathogenic variants

We identified a list of genes that we deemed clinically relevant for reporting cancer-associated germline variants and secondary findings. This list included all cancer risk genes from the BROCA panel (http://tests.labmed.washington.edu/BROCA), as well as the 56 genes from the ACMG guidelines ^5^ for a total of 88 genes (**Supplementary Table S2**).

We devised a variant filtering strategy to narrow down the most important and likely clinically relevant variants (Fig 2). For each variant we collected annotations from database including ClinVar and Exome Aggregation Consortium (ExAC, http://exac.broadinstitute.org) ^20^. Following the ACMG Standards and Guidelines regarding classification and interpretation of sequence variants ^21^, the variants are classified into five categories: Pathogenic Likely Pathogenic, Likely Benign, Benign and variants of unknown significance (VUS) (see Methods for details). A board-certified clinical geneticist reviews all pathogenic and likely pathogenic variants and removes likely misannotated VUS (a rare event). The remaining variants are reported using a detailed report we generate for each case; the other variant categories (Likely Benign, Benign, VUS) are shown in the Appendix of the report we generate for each case.

**Figure 2:**
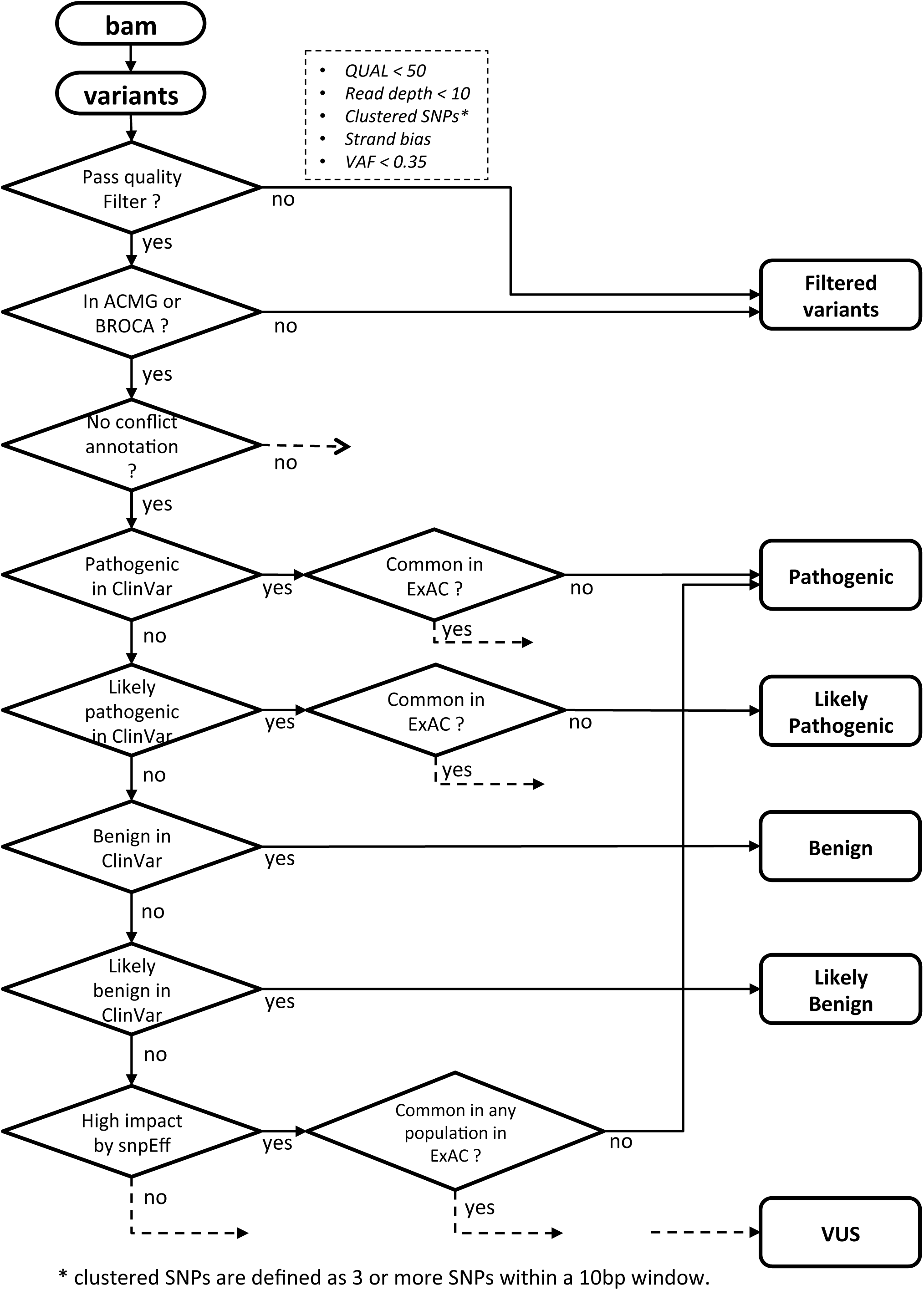
Workflow for screening variants to report. Starting from BAM files, our germline pipeline calls variants using GATK. Variants are annotated using snpEff and snpSift, and then go through a set of filtering schemes: Variants that are 1) having quality < 50; 2) having read depth < 10; 3) clustered; 4) having strand bias; or 5) having variant allele frequency < 0.35; 6) not in our 88 ACMG+BROCA gene panel, are discarded. The remaining variants are classified into five categories: Pathogenic, Likely pathogenic, Likely benign, Benign, and VUS (Variant of unknown significance). Dashed arrow indicates that a variant is assigned to category VUS.

The systematic application of this analytical pipeline to the 343 WES specimens in our cohort identified 60 variants as pathogenic or likely pathogenic, 51 of which were unique. 17% of our patients had a reportable variant on average. We applied the same analysis and filtering strategy to three control cohorts: (a) 2,504 samples from 1000 Genomes Project (1000G) ^22^, (b) 9,282 samples from a WES study on autism (SSC) ^23^, and (c) 128 samples from The Ashkenazi Genome Consortium (TAGC) ^24^. The numbers of variants in total and per individual on average are shown in Table 1; the fully annotated variant lists are in **Supplementary Table S3 (1000G), Supplementary Table S4 (SSC)** and **Supplementary Table S5 (TAGC)**.

**Table 1:** summary of variant calls on the IPM cohort, a subset of IPM cohort after removing Ashkenazi patients, and three control cohorts

| cohort | # samples | # variants | # unique variants | # variants per sample |
| --- | --- | --- | --- | --- |
| IPM | 343 | 60 | 51 | 0.17 |
| IPM_noASH | 241 | 45 | 44 | 0.19 |
| 1000Genomes | 2504 | 370 | 189 | 0.15 |
| SSC | 9282 | 1002 | 365 | 0.11 |
| TAGC | 128 | 24 | 10 | 0.19 |

**Table 2:** table of <u>cancer</u> germline variants found in IPM cohort. The "chr” and "pos” columns record the genomic location of a variant; the "ref” and "alt” columns record the reference and alternative allele of a variant; the "ID” column records the corresponding dbSNP id, ".” indicates no dbSNP ids were found; the column "effect” indicates the effect of a variant (e.g. nonsynonymous, frame shift, etc); the "gene” column records the gene name; the "DNA change” and "AA change” columns record the changes in coding sequence and the changes in amino acid sequence; the "transcript” column records the corresponding Ensembl id of the affected transcript; the "ACMG” and "BROCA” columns indicate whether or not a variant is on ACMG or BROCA genes; the "Count” and "Frequency” columns indicate the number of samples carrying a variant and the corresponding percentage in our IPM cohort; the "site” column records the primary tumor sites of patients carrying a given variant.

| chr | pos | ref | alt | ID | effect | gene | DNA change | AA change | transcript | ACMG | BROCA | Count | Frequency | site |
| --- | --- | --- | --- | --- | --- | --- | --- | --- | --- | --- | --- | --- | --- | --- |
| 5 | 112175211 | T | A | rs1801155 | NON_SYNONYMOUS | APC | c.3920T>A | p.Ile1307Lys | ENST00000257430 | yes | yes | 5 | 1.46% | Prostate; Bladder; Kidney |
| 17 | 7578457 | C | T | . | NON_SYNONYMOUS | TP53 | c.473G>A | p.Arg158His | ENST00000269305 | yes | yes | 2 | 0.58% | Retroperitoneum; Brain |
| 16 | 15844048 | G | A | rs111404182 | NON_SYNONYMOUS | MYH11 | c.2026C>T | p.Arg676Cys | ENST00000396324 | yes | no | 2 | 0.58% | Esophagus; Soft Tissue |
| 13 | 32914437 | GT | G | rs80359550 | FRAME_SHIFT | BRCA2 | c.5946delT | p.Ser1982fs | ENST00000380152 | yes | yes | 2 | 0.58% | Prostate |
| 13 | 32915082 | CTG | C | rs80359606 | FRAME_SHIFT | BRCA2 | c.6591_6592delTG | p.Glu2198fs | ENST00000380152 | yes | yes | 1 | 0.29% | Prostate |
| 15 | 67391213 | TG | T | . | FRAME_SHIFT | SMAD3 | c.5delG | p.Gly2fs | ENST00000559092 | yes | no | 1 | 0.29% | Brain |
| 11 | 108203492 | C | T | rs138941496 | STOP_GAINED | ATM | c.7792C>T | p.Arg2598* | ENST00000278616 | no | yes | 1 | 0.29% | N/A |
| 13 | 32911297 | TAAAC | T | rs80359351 | FRAME_SHIFT | BRCA2 | c.2808_2811delACAA | p.Ala938fs | ENST00000380152 | yes | yes | 1 | 0.29% | Prostate |
| 13 | 32903604 | CTG | C | rs80359604 | FRAME_SHIFT | BRCA2 | c.658_659delGT | p.Val220fs | ENST00000380152 | yes | yes | 1 | 0.29% | Prostate |
| 11 | 64572643 | G | A | . | STOP_GAINED | MEN1 | c.1228C>T | p.Gln410* | ENST00000337652 | yes | yes | 1 | 0.29% | Lung |
| 17 | 41243788 | TAGAC | T | rs80357687 | FRAME_SHIFT | BRCA1 | c.3756_3759delGTCT | p.Ser1253fs | ENST00000471181 | yes | yes | 1 | 0.29% | Brain |
| 11 | 108121531 | C | T | . | STOP_GAINED | ATM | c.1339C>T | p.Arg447* | ENST00000278616 | no | yes | 1 | 0.29% | Prostate |
| 22 | 29115433 | AACTG | A | . | FRAME_SHIFT | CHEK2 | c.758_761delCAGT | p.Ser253fs | ENST00000382580 | no | yes | 1 | 0.29% | Brain |
| 11 | 108164065 | TGATA | T | . | FRAME_SHIFT | ATM | c.4642_4645delGATA | p.Asp1548fs | ENST00000278616 | no | yes | 1 | 0.29% | N/A |
| 3 | 10191581 | C | T | rs28940300 | NON_SYNONYMOUS | VHL | c.574C>T | p.Pro192Ser | ENST00000256474 | yes | yes | 1 | 0.29% | Bladder |
| 17 | 41245492 | C | CA | . | FRAME_SHIFT | BRCA1 | c.2055dupT | p.Glu686fs | ENST00000471181 | yes | yes | 1 | 0.29% | Small Intestine |
| 19 | 1226479 | C | A | . | STOP_GAINED | STK11 | c.395C>A | p.Ser132* | ENST00000586243 | yes | yes | 1 | 0.29% | Bone Marrow |
| 17 | 59878709 | C | G | rs149364097 | NON_SYNONYMOUS | BRIP1 | c.1045G>C | p.Ala349Pro | ENST00000259008 | no | yes | 1 | 0.29% | Soft Tissue |
| 22 | 29121087 | A | G | rs17879961 | NON_SYNONYMOUS | CHEK2 | c.599T>C | p.Ile157Thr | ENST00000348295 | no | yes | 1 | 0.29% | Prostate |
| 5 | 112043415 | A | G | rs189807660 | START_LOST | APC | c.1A>G | p.Met1? | ENST00000507379 | yes | yes | 1 | 0.29% | Liver |
| 11 | 108172389 | C | G | . | STOP_GAINED | ATM | c.5192C>G | p.Ser1731* | ENST00000278616 | no | yes | 1 | 0.29% | Prostate |
| 11 | 108154967 | GT | G | . | FRAME_SHIFT | ATM | c.3764delT | p.Leu1255fs | ENST00000278616 | no | yes | 1 | 0.29% | Prostate |

### Clinically relevant non-cancer germline variants are frequently detected in our advanced cancer cohort

Our list of 88 reportable genes included all the ACMG genes, 32 of which are not cancer genes but instead linked to diseases such as cardiomyopathies, Marfan syndrome and others. The reportable variants in non-cancer genes found in our 343 WES cohort are shown in **Supplementary Table S6**. The most represented gene in that list is SCN5A, followed by LDLR, PCSK9, MYH7, KCNH2, DSP, KCNQ1 and DSG2 (**Supplementary Fig S1a**). Mutations in the SCN5A gene (which codes for a sodium channel) have been linked to Brugada syndrome and congenital long QT syndrome. We also identified patients with germline mutations in DSG2, LDLR and MYH7 (linked to hypertrophic and dilated cardiomyopathies), KCNH2 and KCNQ1 (also linked to congenital long QT syndrome and Cardiac arrhythmia), and FBN1 (linked to Marfan syndrome). Most of these genes are also ranked high in the three control cohorts (1000 Genomes; **Supplementary Fig S1b**, Autism; **Supplementary Fig S1c** and Ashkenazim; **Supplementary Fig S1d**). Ashkenazim represent a significant fraction of our patient cohort and are not strongly represented in the ExAC database, which we use to evaluate rarity of variants. However the most-represented non-cancer genes were overall unchanged after removing Ashkenazi individuals from our cohort (**Supplementary Fig S1e**). Altogether we found that 8.8% of patients in our advanced cancer cohort harbored one or more reportable germline variant in a non-cancer gene. However, we found that this frequency was not statistically different than their frequency in the three control cohorts (11.0% in 1000 Genomes cohort, 6.5% in Autism cohort and 6.3% in Ashkenazim) when all ethnicities were considered together (Fig 3a). Altogether these results indicate that clinically relevant non-cancer germline variants are frequent in our advanced cancer cohort, but not more frequent than expected if individuals were randomly sampled from the general population.

**Figure 3:**
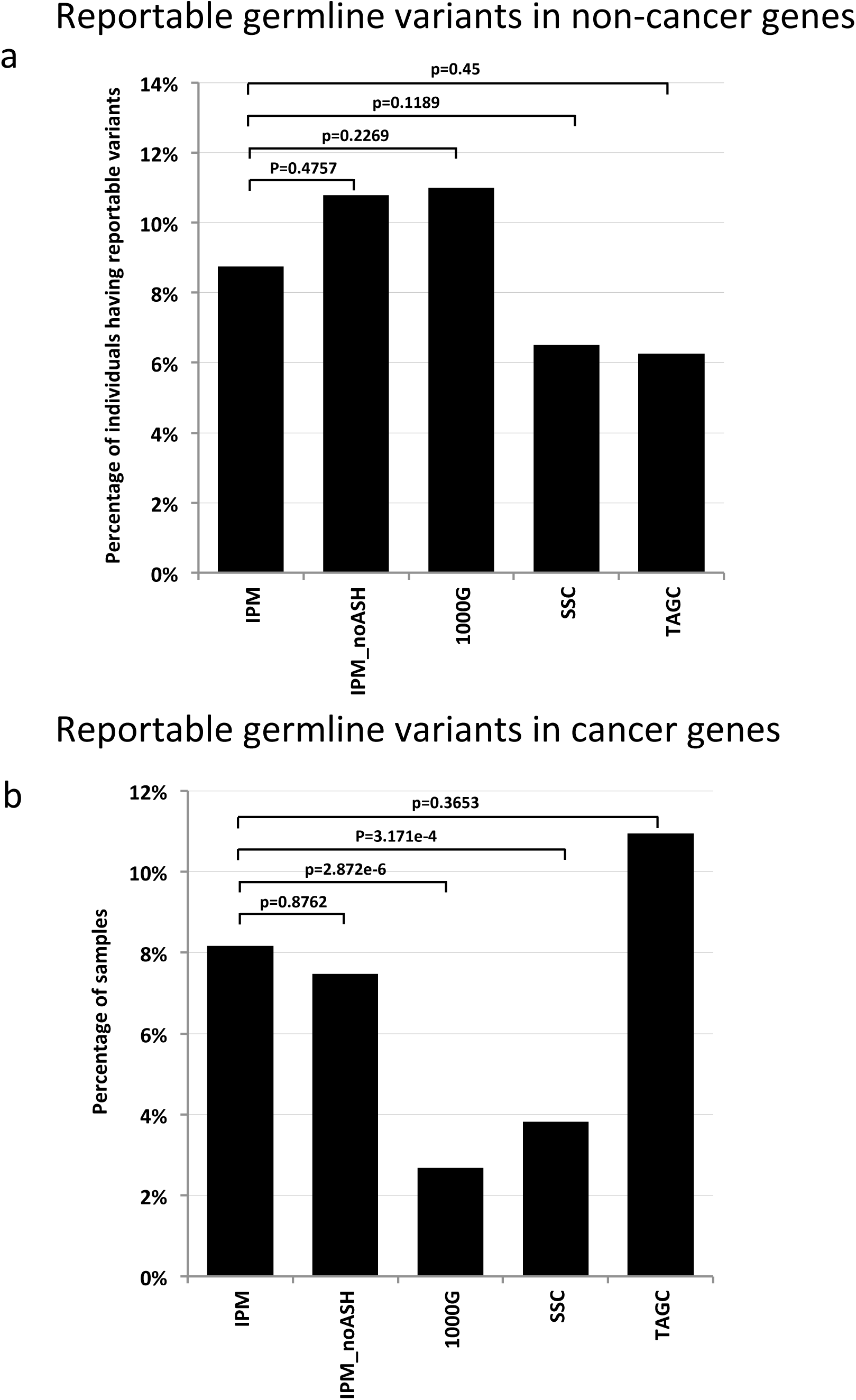
**a) Percentage of patients having reportable germline variants in non-cancer genes across all cohorts.** IPM is our 343-patients cohort. IPM_noASH is a subset of the IPM cohort by excluding Ashkenazi patients. 1000G is the 1000 Genomes cohort. SSC is the Simon's Simplex autism Cohort. TAGC is the Ashkenazi cohort. P-values are calculated using a Fisher's exact test. **b) Percentage of patients having reportable germline variants in noncancer genes across all cohorts.** IPM is our 343-patients cohort. IPM_noASH is a subset of the IPM cohort by excluding Ashkenazi patients. 1000G is the 1000 Genomes cohort. SSC is the Simon's Simplex autism Cohort. TAGC is the Ashkenazi cohort. P-values are calculated using a Fisher's exact test.

### Cancer related disease risk variants are frequent in a cancer precision medicine cohort

We then identified all variants in cancer-associated genes. In total, 56 genes on our list of 88 fall into the cancer category according to the BROCA panel annotation and the Cancer Gene Census from the Sanger Center ^25^. We found that our cohort contains 29 variants within 12 of these genes. The most commonly affected cancer-associated genes in our cohort are APC, BRCA2 and ATM, followed by BRCA1, MYH11, TP53 and CHEK2, with a frequency >0.5%, as shown in **Supplementary Fig S2a**. We found that mutations in certain genes are only prevalent in a specific population. For example, 5 out of the 6 variants found in the APC gene are harbored in Ashkenazi patients; this is in concordance with APC variants being commonly observed in Ashkenazi population with, 8.6% individuals in TAGC cohort carrying variants in this gene (**Supplementary Fig S2b and Fig S2c**). In addition, Ashkenazi patients also contribute two variants found in TP53 gene. As expected from the large size of our two control cohorts (2,504 individuals in 1000G and 9,282 individuals in SSC), we observed a larger absolute number of reportable germline variants in the two non-cancer cohorts (SSC and 1000G) compared to our cohort (Table 1). However unlike in our advanced cancer patient cohort, the majority of these genes were mutated at low frequency (<0.1%) in the two controls cohorts (**Supplementary Fig S2d** and **Supplementary Fig S2e**).

**Table 3** shows the germline variants found in our cohort sorted according to their frequencies. Five distinct patients (three prostate cancer, one bladder cancer and one kidney cancer) were found to harbor a p.I1307K variant in the APC gene, frequent in Ashkenazi individuals (8.6%, 11 out of 128 in the TAGC cohort), linked to an increased risk of colon cancer ^26^ and to several other malignancies ^27^. It has been suggested that Ashkenazi individuals with a p.I1307K variant in APC should be considered for screening colonoscopy at the age of 40, to be repeated every 5 years (similar to recommendations for individuals with family history of colorectal cancer) ^26^. Seven patients had frameshift indels in BRCA1 or BRCA2 genes (2 in BRCA1 and 5 in BRCA2); five of them are known deleterious mutations as annotated by ClinVar. Interestingly, none of these patients were breast or ovarian cancer patients, however this non-association is statistically insignificant due to the small number of breast/ovarian cancer patients in our cohort (n=20). Instead, of the 7 patients with BRCA1 and BRCA2 mutations, 5 males had metastatic prostate cancer, one had neuroendocrine tumor of the small intestine and the last one had astrocytoma. Importantly, 6 out of 7 of these BRCA mutation positive patients had a family history of cancer and in 5 of these cases at least two other family members had received a cancer diagnosis. Five patients had mutations in ATM gene (2 frameshift deletions and 3 nonsense variants), three of them were prostate cancer patients and the other two had unclassified malignancies. Two of the prostate cancer patients with ATM nonsense variants had a family history of cancer (but not prostate cancer). Our findings in metastatic prostate cancer are compatible with the recently reported high incidence of germline variants in genes mediating DNA-repair processes ^6^. Like BRCA1/2, ATM mutations are associated with risk of cancer development, and might be targeted by drugs such as platinum and PARP inhibitors. Two distinct patients were found carrying a missense mutation (p.R158H) in the TP53 gene; both were Ashkenazim and had a family history of cancer. Pathogenic TP53 germline variants are associated with Li-Fraumeni syndrome (LFS), a syndrome associated with a variety of early-onset tumors; it is presently unknown whether these two patients were affected with LFS. For the most part, patients in the cohort had a single reportable cancer-associated variant. However, one patient had both a pathogenic variant in BRCA2 (frameshift deletion) and a p.I1307K variant in APC. Altogether, chart reviews showed that 61% of patients with reportable cancer-associated germline variants had a family history of cancer.

We sought to determine whether the number of cancer-associated variants in this advanced cohort was higher than expected based on numbers obtained in non-cancer cohorts. We found that 8.2% of our patients had cancer-associated variants (Fig 3b), significantly higher than in the 1000 Genomes cohort (2.7%; p=2.9e-6, Fisher's exact test; Fig 3b) and in the autism cohort (3.8%, p=3.2e-4, Fisher's exact test; Fig 3b). The result was confirmed (p=2.8e-4 and p=1.0e-2, respectively) upon removal of the Ashkenazi patients expected to have high number of cancer-associated variants based on the TAGC Ashkenazi cohort (up to 10.9%; Fig 3b). Focusing only on prostate cancer patients in our cohort, we found that up to 12.4% of them (12 out of 97) had reportable cancer-associated variants. This elevated frequency confirms findings from a recent SU2C/PCF prostate cancer precision medicine study that shows high prevalence of germline cancer risk events in advanced prostate cancer patients ^6,28^. In addition, 9.8% (4 out of 41) of patients with brain cancer and 4.8% (2 out of 42) of patients with bladder cancer (the second and third most prevalent cancer type in our cohort, respectively) have reportable cancer-associated variants. The high prevalence of cancer-associated variants in our advanced cancer cohort supports a causal role for these variants in developing their disease and possibly a role in facilitating or accelerating disease progression.

## Discussion

In this study, we presented a preliminary analysis of reportable germline variants in an advanced cancer cohort having undergone tumor-normal whole exome sequencing analysis to identify somatic mutations in tumor biopsies. We observed an increased frequency of cancer-associated reportable germline variants in our advanced cancer cohort compared to two noncancer control cohorts (1000 Genomes cohort and autism cohort). Confirming other observations ^6,28–30^ we observe that variants in known cancer risk genes such as BRCA1 and BRCA2 are found in patients with tumor type that extend beyond ovarian and breast cancer. Review of the patient charts revealed that 61% of patients with reportable variants in cancer genes had a family history of cancer. This was particularly acute in BRCA mutation positive patients, where 6 out of 7 of these had a family history of cancer and in 5 of these cases at least two other family members had received a cancer diagnosis. These results together with previous ones ^28^ indicate that germline risk testing may need to be extended beyond the more commonly recommended breast and ovarian cancer and perhaps systematically offered in patients with any family history of cancer. In addition, these results suggest that when such variants are encountered in the process of somatic tumor testing (as was performed in the present study), they should likely be reported to treating physicians and patients, provided that appropriate consenting and genetic counseling is in place. In some cases such as BRCA gene loss of function mutations, such variants may directly impact clinical cancer care via the use of PARP inhibitors. The immediate need to return non-cancer germline variants in cancer patients is less obvious, especially when tested patients have no clear sign of disease associated with those variants e.g. cardiomyopathies. However, following the ACMG recommendations we currently report all pathogenic variants using the report structure shown in **Supplementary Fig S3**.

One key question in a cancer cohort such as ours is whether there are any correlations between germline and somatic variants. In one model, germline variants act as inactivating (or activating) hit of the first allele and somatic variants act as second hit in the same gene. When we looked for secondary hits (point mutations or indels) in the germline variants in our 56 cancer genes, we found no such events beyond the FANCA gene example that we already reported as relevant to cisplatin sensitivity ^15^. We cannot however exclude the possibility that a secondary hit resides in other cancer genes, as well as other mechanisms acting as secondary hits, such as copy number losses or gains or DNA methylation changes.

Our current strategy for identifying and reporting secondary germline findings can be expanded in a number of directions. An important direction is to implement a more systematic ethnicity-based annotation. Some variants can be rare when assessed across population but may turn common in one particular population. For example, the APC p.I1307K variant is rare in the ExAC database (0.17%) but is commonly observed in the TAGC cohort (8.6%) (**Supplementary Table S7 and S8**). Increasing access to large ethnically diverse cohorts, such as the Precision Medicine Initiative / All of Us cohort ^31^ will likely further facilitate interpretation of such variants using the appropriate ethnicity context. A third direction is to build a well-curated germline variant knowledge database. Our current variant annotations are mainly based on the ClinVar database in which a variant may have multiple annotations submitted from different research/clinical labs. In some cases annotations on the same variant can be inconsistent. As a result, the true significance of such variants may be unclear. In addition, annotations in ClinVar are limited, giving only a score that indicates clinical significance and a short description about the related disease; sometimes the disease name is unknown. It may soon become highly desirable to build a more comprehensive, clinically oriented germline knowledge base that will include more information about germline variant. We have recently released the Precision Medicine Knowledge Base ^32^, a database for interpreting clinically relevant somatic cancer variants. Such databases can be extended to cover germline variants. One more direction for improvement involves germline copy number variants (CNV), currently not called out by our pipeline. A recent study by our group has discovered two germline CNVs that are strongly associated with prostate cancer ^33^. This underscores the potential usefulness of including CNV in germline analyses related to cancer. While calling CNVs is notoriously difficult with whole-exome sequencing, a number of tools introduced in recent years may enable accurate CNV calling e.g., CONTRA ^34^, XHMM ^35^, and CoNIFER ^36^. Integrating these tools in our pipeline together with categorization of CNV (both evidence-based and clinical) represents an important direction for further research.

## Methods

### Whole exome sequencing and analytical pipeline for detecting and prioritizing germline variants

Germline samples used in this study were either blood or buccal swab or normal tissue from formalin-fixed paraffin embedded specimens or frozen tissue (see ^15,16^ for further details). All samples were sequenced to serve as matched controls in the context of CLIA/CLEP approved genomic testing to identify somatic mutations in tumors. As indicated in ^15,16^, exome capture was performed with the Agilent HaloPlex Kit. Paired-end sequencing (2 × 100 bp) was performed on the Illumina HiSeq2500 instrument. The sequenced reads were cleaned by trimming adapter sequences and low quality bases, and were aligned to the human GRCh37 reference using Burrows-Wheeler Aligner (BWA). The aligned bam file was further refined by indel realignment and base quality recalibration with the Genome Analysis Toolkit (GATK). We then used GATK UnifiedGenotyper to call single nucleotide variants (SNVs) and indels, and filtered out variants that are: 1) of low quality (QUAL score < 50); 2) with less than 10 supporting reads; 3) clustered (defined as 3 or more SNVs within a 10bp window); 4) with strand bias; 5) variant allele frequency less than 0.35. Specificity was calculated as TN/(TN+FP); sensitivity as TP/(TP+FN); false discovery rate as TP/(FP+TP) where TN is number of true negative calls, TP is number of true positive calls, FP is number of false positive calls, FN is number of false negative calls. We used the SnpEff and SnpSift packages to generate annotations (for example, gene location, amino acid changes, etc) on the called variants, to predict the impact of a variant on a given gene, and to integrate annotations from ClinVar (version 20160531) and ExAC database (version r0.3.1). We filtered out variants that were: 1) not in the 88 ACMG/BROCA gene list; 2) having unclear disease name in ClinVar; 3) having conflict annotations in ClinVar; 4) heterozygous variants on the MUTYH gene (since this gene is autosomal recessive as annotated by ACMG). The remaining variants were categorized into five categories for reporting: ***Pathogenic*** - variants that are either annotated as pathogenic in the ClinVar database and have allele frequency less than 0.01 as annotated by ExAC, or predicted as HIGH impact by SnpEff and have allele frequency less than 0.01 in any populations in ExAC; ***Likely Pathogenic*** - variants that are annotated as likely pathogenic in ClinVar and have allele frequency less than 0.01 in ExAC; ***Likely benign*** - variants that are annotated as likely benign in ClinVar; ***Benign*** - variants that are annotated as benign in ClinVar; ***VUS*** - variants that cannot be assigned to the above four categories and are of unknown significance.

### Ethnicity inference

Ethnicity analysis was performed by means of Ethseq (Romanel et. al, submitted), a computational tool that infers the ethnicity of a set of individuals based on differential SNPs genotype profiles. Briefly, first, by combining genotype data of individuals with known ethnicity, an NGS platform-specific *reference model* is built. Then, the tool infers the genotype of all considered SNPs for all individuals of interest using ASEQ ^37^ and builds the *target model.* Principal component analysis (PCA) is then performed using the *smartpca* module ^38^ on aggregated target and reference models genotype data.. Euclidian space defined by the first two PCA components is then inspected to, first, generate the smallest convex sets identifying the four main ethic groups (EUR, AFR, EAS and SAS) and then to annotate the ethnicity of the individuals of interest. Individuals lying inside a ethnic group set are annotated with the corresponding ethnicity and INSIDE label; individuals lying outside ethnic group sets are annotate with the nearest (Euclidean distance) ethnic group and CLOSEST label. In this study, we used the 1,000 Genome Project data for the reference model comprising 91,661 SNPs within the WES HaloPlex captured regions. To more precisely define the fraction of Ashkenazi population in our study cohort and to better discern between the spatially close ASH and EUR groups, we implemented a further analysis step by first extending the reference model with Ashkenazi genome data ^24^ and then, by reducing both the reference and the target models to EUR and ASH individuals only; for the target model only individuals annotated as INSIDE in the first analysis step were considered.

## Acknowledgments

This work was partially supported by NSF CAREER, LLS SCOR, Hirschl Trust Award, Starr Cancer Consortium I6-A618, NIH 1R01CA194547 to O.E., Department of Defense PC121341, Damon Runyon Cancer Research Foundation-Gordon Family Clinical Investigator Award CI-67- 13 to H.B., Starr Cancer Consortium I7-A771 to H.B. and M.A.R., European grant from the Nuovo-Soldati Foundation to J.C., Italian Association for Cancer Research (AIRC to F.D.), a Simons Foundation Autism Research Initiative grant to 1.1. (SF362665), as well as a Medical Scientist Training Program grant from the National Institute of General Medical Sciences of the National Institutes of Health under award number T32GM007739 to the Weill Cornell/Rockefeller/Sloan Kettering Tri-Institutional MD-PhD Program for Y.X. We also acknowledge generous financial support from Caryl and Israel Englander to the Institute for Precision Medicine. We are grateful to Mike Wigler for his suggestion to use the SSC as control cohort. We also thank Michael Dorschner and Gail Jarvik for helpful discussions and suggestions.

## References

1. Kiezun A, Garimella K, Do R, et al. Exome sequencing and the genetic basis of complex traits. Nat Genet 2012;44(6):623–630.

2. Van Allen EM, Wagle N, Stojanov P, et al. Whole-exome sequencing and clinical interpretation of formalin-fixed, paraffin-embedded tumor samples to guide precision cancer medicine. Nat Med 2014;20(6):682–688.

3. Beltran H, Eng K, Mosquera JM, et al. Whole-Exome Sequencing of Metastatic Cancer and Biomarkers of Treatment Response. JAMA Oncol 2015;1(4):466–474.

4. Roychowdhury S, Iyer MK, Robinson DR, et al. Personalized oncology through integrative high-throughput sequencing: a pilot study. Science translational medicine. 2011;3(111):111ra121.

5. Green RC, Berg JS, Grody WW, et al. ACMG recommendations for reporting of incidental findings in clinical exome and genome sequencing. Genetics in medicine : official journal of the American College of Medical Genetics 2013;15(7):565–574.

6. Pritchard CC, Mateo J, Walsh MF, et al. Inherited DNA-Repair Gene Mutations in Men with Metastatic Prostate Cancer. N Engl J Med 2016;375(5):443–453.

7. Thorn CF, Klein TE, Altman RB. PharmGKB: the Pharmacogenomics Knowledge Base. Methods Mol Biol 2013;1015:311–320.

8. Directors ABo. ACMG policy statement: updated recommendations regarding analysis and reporting of secondary findings in clinical genome-scale sequencing. Genetics in medicine : official journal of the American College of Medical Genetics. 2015;17(1):68–69.

9. Tabor HK, Auer PL, Jamal SM, et al. Pathogenic variants for Mendelian and complex traits in exomes of 6,517 European and African Americans: implications for the return of incidental results. Am J Hum Genet 2014;95(2):183–193.

10. Dorschner MO, Amendola LM, Turner EH, et al. Actionable, pathogenic incidental findings in 1,000 participants' exomes. Am J Hum Genet 2013;93(4):631–640.

11. Amendola LM, Dorschner MO, Robertson PD, et al. Actionable exomic incidental findings in 6503 participants: challenges of variant classification. Genome Res. 2015.

12. Ng D, Johnston JJ, Teer JK, et al. Interpreting secondary cardiac disease variants in an exome cohort. Circulation Cardiovascular genetics 2013;6(4):337–346.

13. Jurgens J, Ling H, Hetrick K, et al. Assessment of incidental findings in 232 whole-exome sequences from the Baylor-Hopkins Center for Mendelian Genomics. Genetics in medicine : official journal of the American College of Medical Genetics. 2015.

14. Zhang J, Walsh MF, Wu G, et al. Germline Mutations in Predisposition Genes in Pediatric Cancer. N Engl J Med 2015;373(24):2336–2346.

15. Beltran H, Eng K, Mosquera JM, et al. Whole Exome Sequencing of Metastatic Cancer and Biomarkers of Treatment Response. JAMA Oncology 2015;1(4):466–474.

16. Rennert H, Eng K, Zhang T, et al. Development and validation of a whole-exome sequencing test for simultaneous detection of point mutations, indels and copy-number alterations for precision cancer care. npj Genomic Medicine. 2016;1:16019.

17. Li H, Durbin R. Fast and accurate short read alignment with Burrows-Wheeler transform. Bioinformatics 2009;25(14):1754–1760.

18. DePristo MA, Banks E, Poplin R, et al. A framework for variation discovery and genotyping using next-generation DNA sequencing data. Nat Genet. 2011;43(5):491–498.

19. Zook JM, Chapman B, Wang J, et al. Integrating human sequence data sets provides a resource of benchmark SNP and indel genotype calls. Nat Biotechnol. 2014;32(3):246–251.

20. Lek M, Karczewski KJ, Minikel EV, et al. Analysis of protein-coding genetic variation in 60,706 humans. Nature 2016;536(7616):285–291.

21. Richards S, Aziz N, Bale S, et al. Standards and guidelines for the interpretation of sequence variants: a joint consensus recommendation of the American College of Medical Genetics and Genomics and the Association for Molecular Pathology. Genetics in medicine : official journal of the American College of Medical Genetics 2015;17(5):405–424.

22. Genomes Project C, Abecasis GR, Auton A, et al. An integrated map of genetic variation from 1,092 human genomes. Nature 2012;491(7422):56–65.

23. Iossifov I, O'Roak BJ, Sanders SJ, et al. The contribution of de novo coding mutations to autism spectrum disorder. Nature 2014;515(7526):216–221.

24. Carmi S, Hui KY, Kochav E, et al. Sequencing an Ashkenazi reference panel supports population-targeted personal genomics and illuminates Jewish and European origins. Nat Commun 2014;5:4835.

25. Futreal PA, Coin L, Marshall M, et al. A census of human cancer genes. Nat Rev Cancer 2004;4(3):177–183.

26. Boursi B, Sella T, Liberman E, et al. The APC p.I1307K polymorphism is a significant risk factor for CRC in average risk Ashkenazi Jews. European journal of cancer 2013;49(17):3680–3685.

27. Leshno A, Shapira S, Liberman E, et al. The APC I1307K allele conveys a significant increased risk for cancer. Int J Cancer 2016;138(6):1361–1367.

28. Robinson D, Van Allen EM, Wu YM, et al. Integrative clinical genomics of advanced prostate cancer. Cell 2015;161(5):1215–1228.

29. Ferrone CR, Levine DA, Tang LH, et al. BRCA germline mutations in Jewish patients with pancreatic adenocarcinoma. J Clin Oncol 2009;27(3):433–438.

30. Levy-Lahad E, Friedman E. Cancer risks among BRCA1 and BRCA2 mutation carriers. Br J Cancer 2007;96(1):11–15.

31. Collins FS, Varmus H. A new initiative on precision medicine. N Engl J Med 2015;372(9):793–795.

32. Huang L, Fernandes H, Zia H, et al. The cancer precision medicine knowledge base for structured clinical-grade mutations and interpretations. J Am Med Inform Assoc. 2016.

33. Demichelis F, Setlur SR, Banerjee S, et al. Identification of functionally active, low frequency copy number variants at 15q21.3 and 12q21.31 associated with prostate cancer risk. Proc Natl Acad Sci U S A 2012;109(17):6686–6691.

34. Li J, Lupat R, Amarasinghe KC, et al. CONTRA: copy number analysis for targeted resequencing. Bioinformatics 2012;28(10):1307–1313.

35. Fromer M, Moran JL, Chambert K, et al. Discovery and statistical genotyping of copy-number variation from whole-exome sequencing depth. Am J Hum Genet 2012;91(4):597–607.

36. Krumm N, Sudmant PH, Ko A, et al. Copy number variation detection and genotyping from exome sequence data. Genome Res. 2012;22(8):1525–1532.

37. Romanel A, Lago S, Prandi D, Sboner A, Demichelis F. ASEQ: fast allele-specific studies from next-generation sequencing data. BMC Med Genomics 2015;8:9.

38. Price AL, Patterson NJ, Plenge RM, Weinblatt ME, Shadick NA, Reich D. Principal components analysis corrects for stratification in genome-wide association studies. Nat Genet 2006;38(8):904–909.

